# *In vivo* virus-free lineage tracing and live imaging reveal that NeuroD1 does not reprogram microglia into neurons

**DOI:** 10.64898/2026.06.08.730780

**Authors:** Xiaoyu Li, Yuxin Li, Yue Cao, Niewen Hu, Baozhi Yang, Pei Ouyang, Yuxiao Jin, Shuai Gao, Li Cao, Bo Peng, Yanxia Rao

## Abstract

*In situ* glia-to-neuron conversion induced by a single transcription factor is a promising regenerative strategy for treating central nervous system injuries and neurodegenerative diseases. Previous studies reported that NeuroD1 can induce microglia-to-neuron cross-lineage conversion and improve recovery after brain injury. However, these findings remain controversial. To examine the ability of NeuroD1 in inducing microglia-to- neurons conversion, we employed virus-free genetic lineage tracing systems to specifically induce NeuroD1 expression in microglia, and track the cell fate of NeuroD1-expressing cells at multiple timepoints. Meanwhile, we longitudinally monitored NeuroD1-expressing microglia via two-photon imaging, and characterized their transcriptional profile by single-cell RNA sequencing. All observations across these methods revealed that NeuroD1-expressing cells retained their microglia identity, and cannot convert into neurons, no matter under physical and injury conditions. Instead, sustained expression of NeuroD1 facilitated microglia apoptosis *in vivo*. Collectively, our findings provide strong evidence that NeuroD1 alone is sufficient to induce microglia-to neuron conversion. This study further highlighted the importance of rigorous lineage-tracing and cell fate mapping strategies for validating in situ glia-to-neuron conversion.

## Introduction

In the central nervous system (CNS), neurons have limited regenerative capacity, therefore, brain injury and neurodegeneration result in permanent neuronal loss. In contrast, glia cells, especially microglia exhibit a high regenerative capacity(1–3). *In situ* reprogramming glial cells into neurons offers a promising strategy for neuronal regeneration and functional recovery in brain injuries and neurodegenerative disease. Recent years, single gene such as NeuroD1 (4), PAX6 (5), SOX2 (6–8), ASCL1 (9) and PTBP1 (10, 11) have been reported to induce inter-neuroectodermal lineage conversion, transforming astrocytes, oligodendrocyte precursor cells (OPCs) or Muller glia into neurons, shedding a light for neurodegenerative therapies.

However, before clinical translation, these approaches require rigorous validation using stringent and robust approaches. Unbaised and well-controlled lineage tracing, and unambiguous live imaging are regard as gold standards for validating glia-to-neuron conversion (12, 13). Indeed, several studies employing rigorous lineage tracing and live imaging techniques have failed to observe some inter-lineage glia-to-neuron conversions with single-gene manipulation (14–22), including NeuroD1 (20–22). In addition to inter-lineage conversion, Nakashima et al. have reported that NeuroD1 can induced microglia-to-neuron cross-lineage conversion (23). We previously conducted lineage tracing using virus-based tracing system, *in vitro* live-cell imaging, and microglia ablation, and demonstrated that the reported cross-lineage conversion of microglia into neurons may be attribute to experimental artifacts, e.g., off-target transduction from virus (13). Although Nakashima et al. argued that the phenomenon we observed was due to insufficient NeuroD1 expression in microglia (24), our comparison of the data from both studies (13, 23) indicates that our NeuroD1 expression levels are comparable to, or even higher than, those reported in their study (25). Besides, Nakashima et al. reported that NeuroD1 could induce the microglia-to neurons conversion at brain injury sites *in vivo*, and promote recovery after brain injury (26). Such conflicting reports are of particular concern, because they directly influence whether glia-to-neuron reprogramming could be translated in to clinical. To settle this controversy, we employed virus-free genetic lineage tracing system to avoid potential experimental artifacts, longitudinal two-photon live imaging to direct monitor the changes in NeuroD1-expressing microglia *in vivo*, and single cell RNA sequencing (scRNA-seq) to characterize their transcriptional profiles. Our findings provide strong evidence that microglia do not undergo conversion into neurons *in vivo,* under either physiological or pathological conditions. Our study highlighted the importance of employing rigorous lineage tracing, longitudinal live cell imaging, and multimode validation in evaluating glia-to neuron reprogramming approaches.

## Results

### Genetic lineage tracing reveals NeuroD1 expression in microglia fails to induce neuron conversion *in vivo*

We previous reported that ectopic expression of NeuroD1 using lentiviral vector does not induce the microglia-to-neurons conversion, but triggers microglial cell death (13). However, most viral tools designed for microglia exhibit limited efficiency and specificity in the brain, but displayed off-target transduction of neurons and astrocytes (13, 20, 27). This limitation was observed in either lentivirus or adeno-associated virus (AAV) (with different serotypes 6, 8 and 9), and irrespective of the use of myeloid CX3CR1 or CD68 promoters (Figure S1). Similar observations have been reported in our previous study and other groups (13, 20, 27).

Direct evidence for glia-to-neuron conversion by NeuroD1 via a virus-free expression system, particularly genetic lineage tracing is lacking. Genetic lineage tracing are essential for avoiding experimental artifacts relate to virus leakage (12, 13). Thus, we utilized transgenic animals to induce NeuroD1 specifically expression in microglia, and track the cell fate of NeuroD1-expressing microglia. We therefore crossed macrophage/microglia-targeting CX3CR1-CreER mouse line, or a microglia-specific TMEM119-CreER mouse line (28, 29), with an inducible NeuroD1-expressing mouse line Rosa26-LoxP-Stop-LoxP-NeuroD1-IRES-GFP (LSL-NeuroD1-GFP) (30), to obtain CX3CR1-CreER::LSL-NeuroD1-GFP mice (hereafter, CX3CR1-ND1) and TMEM119-CreER::LSL-NeuroD1- GFP mice (hereafter, TMEM119-ND1 for short). Meanwhile, CX3CR1-CreER::Ai14 (CX3CR1-Ai14) and TMEM119-CreER::Ai14 (TMEM119-Ai14) mice were used as controls (Figure 1A and Figure S2A). Following tamoxifen treatment, all GFP positive cells were continuously co-expressed with NeuroD1 (Figure 1B-1D and Figure S2B-2C), enabling us to tracing NeuroD1-expressing cells for lineage tracing. Meanwhile, we observed that theses GFP positive cells retained the characteristics morphology of microglia, exhibiting typical highly ramified process and small cell body (Figure 1 and Figure S2).

**Figure 1.**
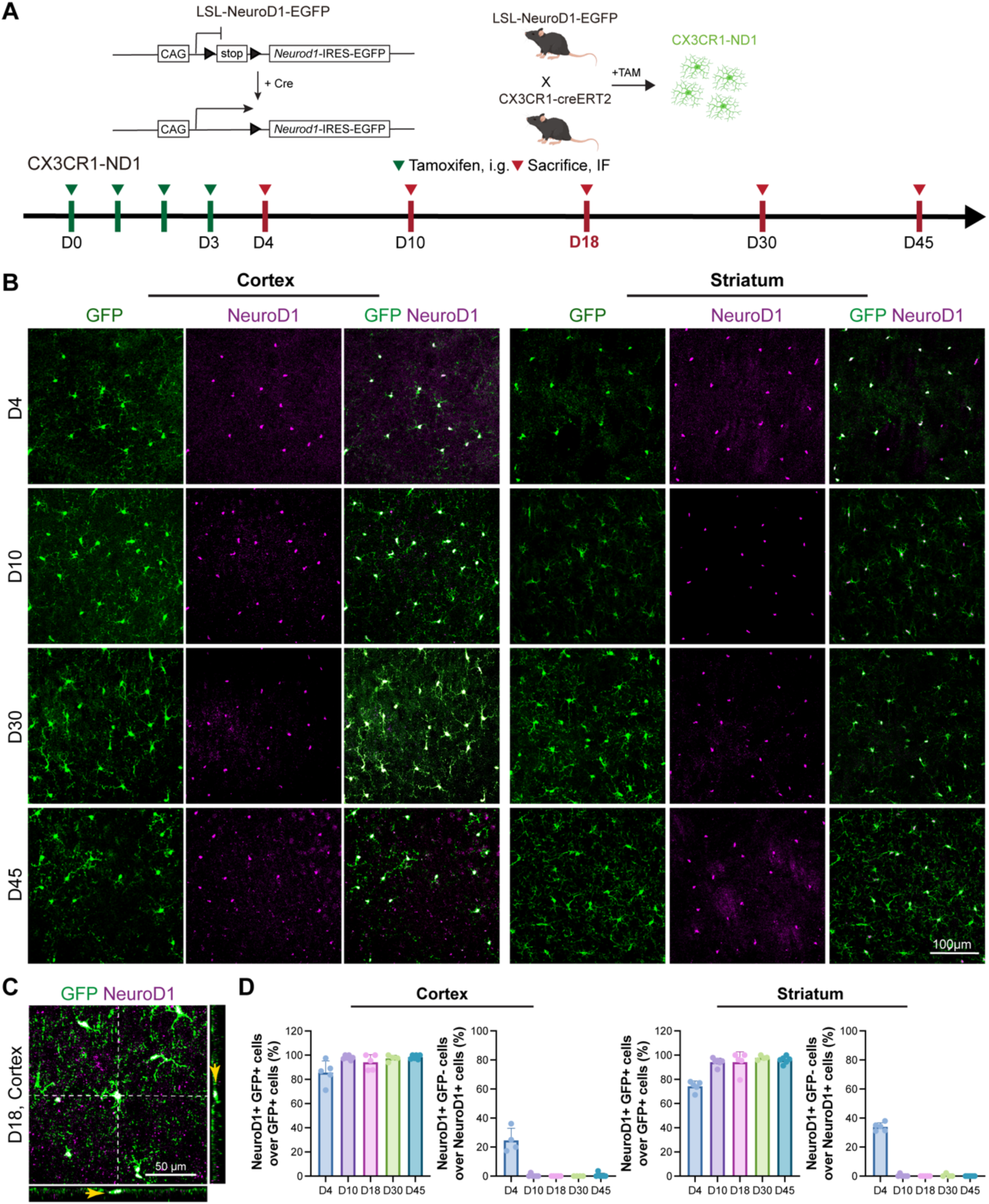
Virus-free genetic induction can successfully induce NeuroD1 expression in microglia. (A) Schematic of the specific expression of NeuroD1, lineage tracing of microglia and experimental timeline. D, day. i.g. oral gavage. (B) Representative confocal images at D4, D10, D30 and D45 showing that NeuroD1 is robustly co-expressed in GFP-positive cells after Cre recombination. (C) Representative confocal images at D18 showing that NeuroD1 is robustly co-expressed in GFP-positive cells after Cre recombination. (D) Quantification of the percentage of NeuroD1^+^GFP^+^ cells among GFP^+^ cells and NeuroD1^+^GFP^-^ cells among NeuroD1^+^, revealing NeuroD1 is highly co-expressed with GFP. N = 4-6 mice for each group. Data are presented as mean ± SD.

We subsequently examined the fate of GFP positive cells at multiple timepoints, at 4, 10, 18, 30, and 45 days post–tamoxifen induction. Immunostaining for microglia and neuron markers revealed that all GFP positive cells were consistently co-localized with microglia marker IBA1, but they did not express neuronal markers, NeuN, TUJ1 and MAP2 within CX3CR1-ND1 mouse brain (Figure 2 and Figure S3A-3D). The observations were the same in TMEM119-ND1 (Figure S2). These findings indicate that sustained NeuroD1 ectopic expression by genetic modification, microglia retain their microglia identify but not acquire a neuronal cell fate.

**Figure 2.**
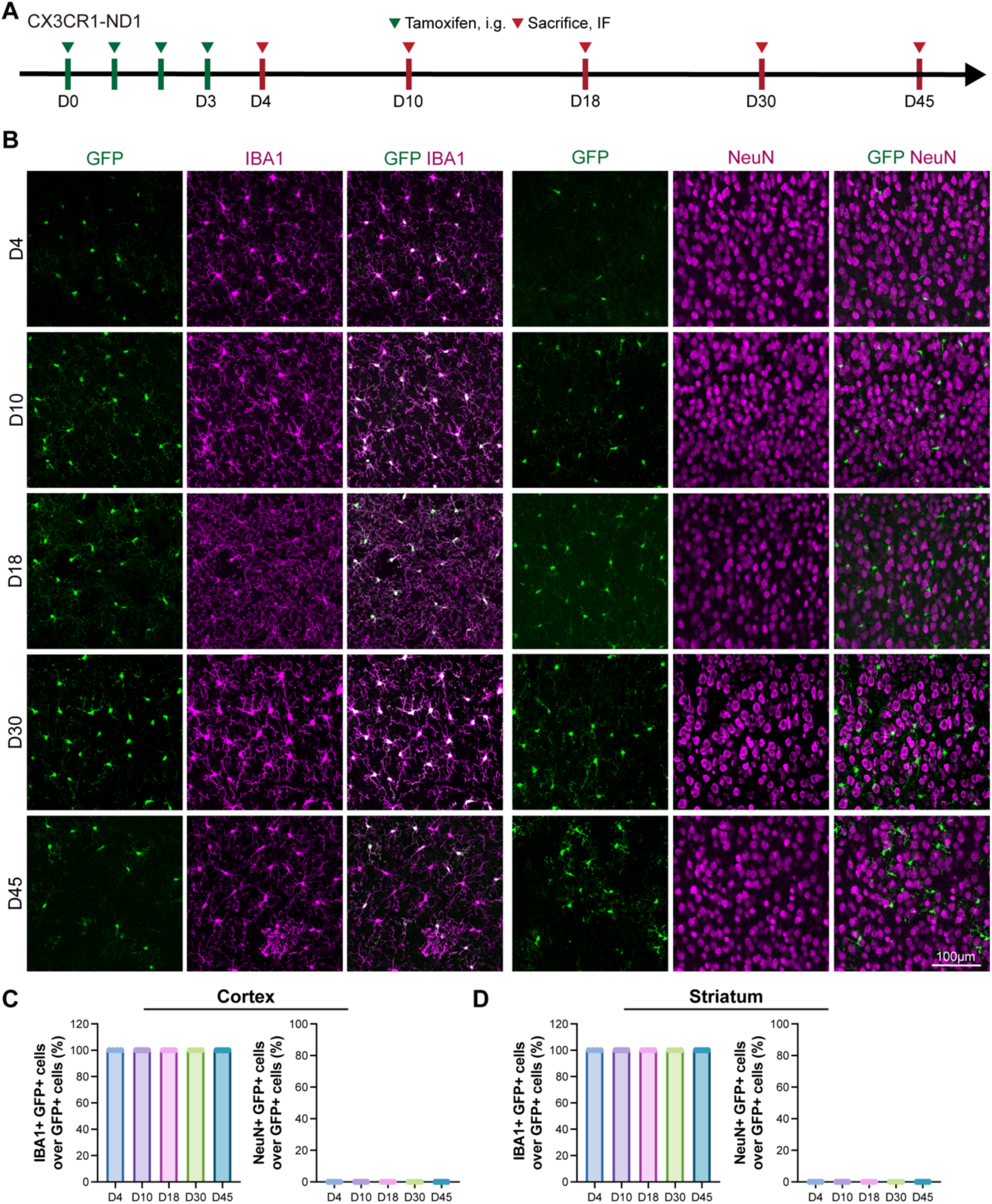
Sustained NeuroD1 expression in microglia does not induce microglia-to-neuron conversion across time. (A) Experimental timeline. D, day. i.g. oral gavage. (B) Representative confocal images at D4, D10, D18, D30 and D45 showing that GFP is co-labeled with IBA1 but not co-labeled with NeuN. (C) Quantification of IBA1^+^GFP^+^ double-positive cells among GFP^+^ cells or NeuN^+^GFP^+^ double-positive cells among NeuN^+^ in cortex at D4, D10, D18, D30 and D45. N = 4 or 5 mice for each group. (D) Quantification of IBA1^+^GFP^+^ double-positive cells among GFP^+^ cells or NeuN^+^GFP^+^ double-positive cells among NeuN^+^ in striatum at D4, D10, D18, D30 and D45. N = 4 or 5 mice for each group.

### Ectopic NeuroD1 expression induce microglia apoptosis rather than neuronal conversion

Given the possibility that genetically labeled cells may loss or downregulation of reporter, such as GFP expression during cell fate conversion (31), we further monitored the cell fate of GFP- labeled microglia *in vivo* using 2-photon microscopy (Figure 3A). We repeated image GFP- positive cells in the frontal cortex with 4 days intervals, beginning on day 18, and continuing throughout day 45 post-tamoxifen administration (Figure 3A). This strategy allowed us to directly track individual NeuroD1-expressing microglia over time. The longitudinal live cell imaging revealed that there was no obvious morphological transition during the imaging period (Figure 3B, white rectangle, and Movie 1). Instead, the GFP-positive cells consistently exhibited a highly ramified morphology with short, and motile processes, which were typical morphological features of microglia rather than neuron (Figure 3B, white rectangle and Movie 1). Meanwhile, we observed that some GFP-labeled microglia progressively disappeared during the imaging period (yellow arrow in Figure 3B, and Movie 1). These observations indicate sustained ectopic NeuroD1 expression could not induce microglia-to-neuron conversion, but induce microglia cell loss *in vivo.* These *in vivo* time lapsing imaging observations were consistent with our previous finding from *in vitro* live cell imaging (13). We further quantified numbers of GFP-labeled microglia at multiple timepoints after tamoxifen induction. The number of GFP-labeled cells initially increased and reached a peak at day 30 post-tamoxifen induction, but subsequently declined by day 45 (Figure 3C, D). The progressive loss of GFP-labeled microglia was also observed in TMEM119-ND1 brain (Figure S2D). These findings further supported that NeuroD1 expression in microglia induce progressively cell loss overtime *in vivo*.

**Figure 3.**
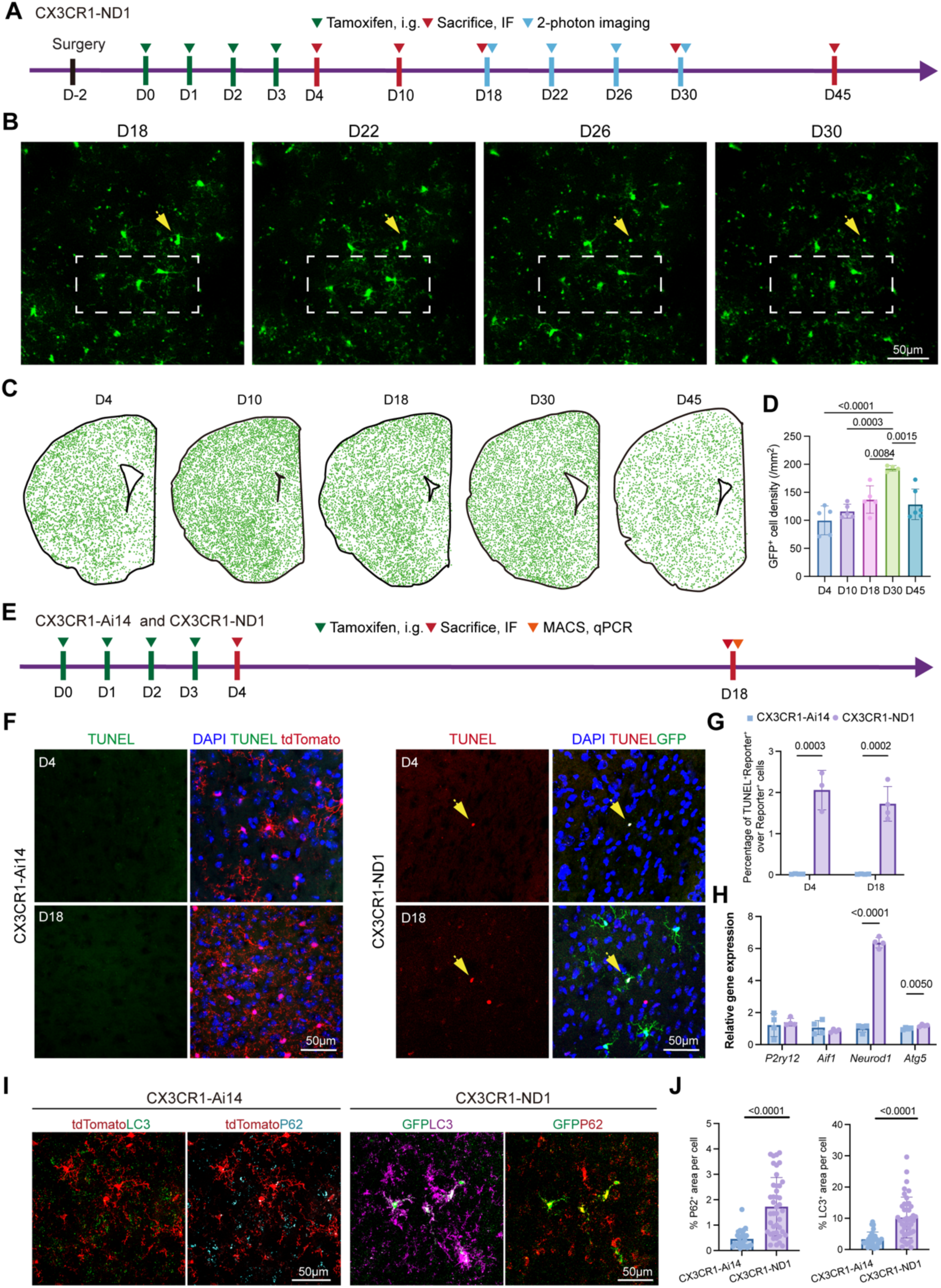
NeuroD1-expressing microglia retain microglial their characteristic morphology, but undergo progressively cell loss over time. (A) Experimental timeline. D, day. i.g. oral gavage. (B) Representative images showing GFP-positive cells with highly ramified morphologies characteristic of microglia rather than neurons. The white rectangular indicate no morphological change, the yellow arrow indicating the GFP^+^ cell loss over time. (C) Representative images illustrating changes in the number of GFP-positive cells over time (green dots indicate GFP-positive cell). (D) Quantification of GFP-positive cells density. N = 4 - 6 mice per time point. Data are presented as mean ± SD. One Way ANOVA with Turkey’s multiple comparison test. (E) Experimental timeline. (F) Representative confocal image shows TUNEL^+^ Reporter^+^ cells are increased in CX3CR1-ND1 animal. (G) Quantification of TUNEL^+^ Reporter^+^ cells at D4 and D18. Unpaired t test. Data are presented as mean ± SD. (H) Relative gene expression level at D18, Unpaired t test. Data are presented as mean ± SD. (I) Representative confocal image shows the LC3 and P62 level in CX3CR1-Ai14 and CX3CR1-ND1mice. Scale bar, 50μm. (J) Quantification of P62^+^ and LC3^+^ area per cell. Unpaired t test. Data are presented as mean ± SD.

To investigate the mechanisms underlying NeuroD1-expressing microglia loss, we first asked that whether the progressive loss of NeuroD1-expressing microglia due to cell apoptosis. To this end, we assessed apoptosis by TUNEL staining in CX3CR1-ND1 and CX3CR1-Ai14 mice (Figure 3E). In line with the gradually disappear of GFP positive cells, we observed that there was a significantly higher proportion of TUNEL positive cells among reporter positive cells in CX3CR1-ND1 mice than that of controls, increasing from approximately 0.01% in controls to 1.7% in NeuroD1-expressing cells (Figure 3F and 3G). This indicate that NeuroD1 expression accelerated microglia apoptosis *in vivo*. We next asked the molecular mechanisms through which NeuroD1 induce microglia cell death. Microglia were isolated from CX3CR1-ND1 and CX3CR1-Ai14 brains using CD11b-based magnetic separation at day 18 post tamoxifen induction (Figure 3E and Figure S3A). We then performed qPCR to examine the expression of genes involved in apoptosis (*Parp1*, *Casp3*, *Bax*, *Bcl2*), necrosis (*Ripk1*, *Fas*, *Mlkl*), autophagy (*Becn1* and *Atg5*) (25, 32, 33). Among the genes examined, *Atg5*, a key regulator of autophagy (34), was significantly upregulated in cells isolated from CX3CR1-ND1 compared with controls (Figure 3H). We therefore examined autophagic markers following NeuroD1 induction. In CX3CR1-ND1 mice, reporter^+^ cells exhibited about 4-fold higher LC3 and P62 levels than controls, as shown by immunostaining (Figure 3I-J), suggesting accumulation of autophagosomes markers and impaired autophagic flux. Taken together, these data indicate that sustained ectopic NeuroD1 expression induces microglia apoptosis, accompanied with autophagic dysregulation, leading to NeuroD1-expressing microglia loss in vivo, rather than microglia-to-neuron conversion.

### NeuroD1 fails to reprogram microglia into neurons even in the injured brain

Previous studies suggest that injury environments might precondition glial cells for fate reprogramming. For example, Heinrich et al. demonstrated that a stab wound prior to virus delivery is necessary to stimulate NG2 glia to neuron conversion by SOX2(35). Similarly, Irie et al. showed that NeuroD1 can directly reprogram microglia/macrophages into neurons at the lesion site following transient middle cerebral artery occlusion(26). These findings raise the possibility that the failure of NeuroD1-mediated microglia-to-neuron conversion under physiological conditions may attribute to the absence of injury-induced reprograming permissive microenvironment.

To test this possibility, we established a moderate traumatic brain injury (TBI) in the motor cortex in CX3CR1-ND1 mice as previously describe (36), followed by tamoxifen induction to activate NeuroD1 expression in microglia (Figure 4A). If the injury environment stimulates NeuroD1-meidiated microglia to neuron conversion, the new born neurons would contribute to motor function recovery after TB1. We therefore performed a serries behavior test to assess motor recovery at 19 days post tamoxifen induction (Figure 4A). In grip strength task, the CX3CR1-ND1 mice showed comparable performance to that of controls, with no significant differences in the strength between groups (Figure 4B). Similarly, no improvements were observed in rotarod task, as reflected by a comparable latency to fall between groups (Figure 4C). These results indicate that NeuroD1 expression in microglia does not improve the motor function recovery in TBI model. We next directly examined the cell fate of NeuroD1-expressing microglia in the injury brain. Immunostaining analysis showed that GFP-labeled NeuroD1-expressing microglia did not co-localize with the neuronal marker NeuN (Figure 4D, E) or MAP2 (Figure 4F, G), both in the lesion core and distal regions. To further figure out the cell fate of microglia after NeuroD1 induction, we conducted scRNA-seq to characterize the transcriptional profile of NeuroD1-expressing cells (Figure 4A). Both tdTomato and GFP positive cells were isolated from CX3CR1-Ai14 and CX3CR1-ND1 brain respectively, using flow cytometry (Figure S4A), under both physiological and TBI condition. These cells were subsequently subjected to scRNA-seq (Figure S4B-4E). In line with the immunofluorescent findings, when we annotated sorted reposter-positve cells, we did not identify any neuronal cluster, even under injury condition (Figure 4H, Figure S4E). Additionally, the reporter-positive cells still strong expressed microglia markers, including *P2ry12*, *Hexb*, and *Tmem119* (Figure 4I and 4J), but did not express neuronal markers, such as *Rbfox3*, *Tubb3*, and *Map2* (Figure 4I and 4J). Collectively, these findings demonstrate that cannot enable NeuroD1 fails to induce microglia-to-neuron conversion, even under an injury condition.

**Figure 4.**
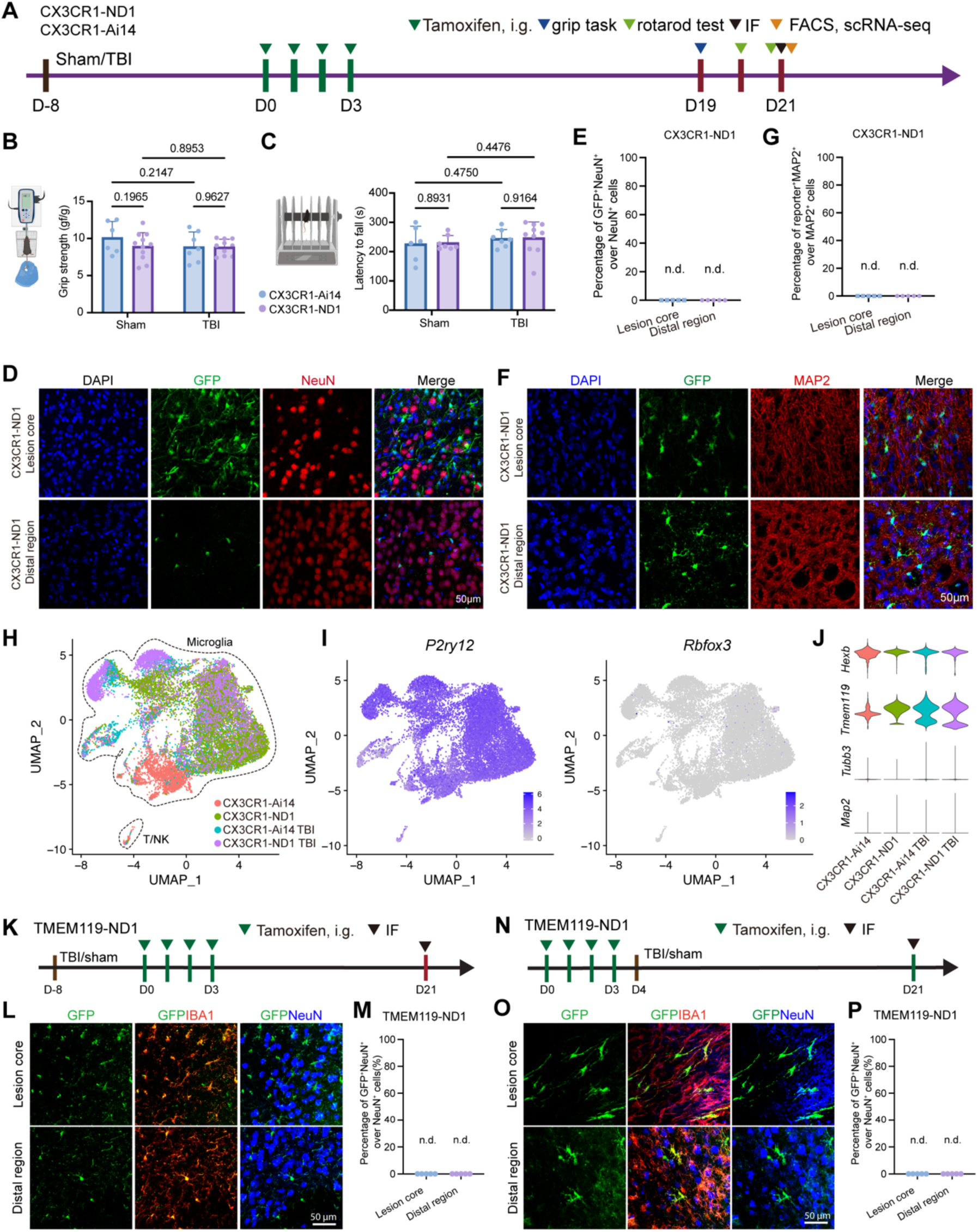
NeuroD1 does not induce microglia-to-neuron conversion even in an injury conditioned microenvironment. (A) Experimental design. D, day. i.g. oral gavage. (B) Quantification of grip strength showing no improvement in NeuroD1-expressing TBI mice. Two-way ANOVA with multiple comparisons. (C) Quantification of rotarod performance showing no improvement in NeuroD1-expressing TBI mice. Two-way ANOVA with multiple comparisons. (D) Representative confocal images of DAPI, GFP and NeuN co-staining results in CX3CR1-ND1 TBI mice, showing no co-localization of GFP with NeuN in either the lesion core or distal regions. Scale bar, 50 μm. (E) Quantification of the percentage of GFP^+^NeuN^+^ cells among total NeuN^+^ cells in both the lesion area and distal area. n.d., not detected. (F) Representative confocal images of DAPI, GFP, and MAP2 in CX3CR1-ND1 mice, showing no co-localization of GFP with MAP2 in either the lesion core or distal regions. Scale bar, 50 μm. (G) Quantification of the percentage of GFP^+^MAP2^+^ cells among total MAP2^+^ cells in the lesion area and distal region. n.d., not detected. (H) UMAP reduction shows the cell types of the FACS-sorted reporter^+^ cells in each group; note that there are no neurons. (I) Feature plot showing that the microglia-specific marker *P2ry12* was highly expressed even under TBI conditions, but the neuron-specific marker *Rbfox3* was barely expressed. (J) Violin plot showing the expression of microglia-specific genes (*Hexb* and *Tmem119*) and neuron-specific genes (*Tubb3* and *Map2*) in 4 groups. (K) Experiment timeline of TMEM119-ND1 mice, TBI was performed before tamoxifen induction. (L) Representative confocal images of GFP, IBA1 and NeuN in TMEM119-ND1 mice, showing no co-localization of GFP with NeuN in either the lesion core or distal regions. Scale bar, 50 μm. (M) Quantification of the percentage of GFP^+^NeuN^+^ cells among total NeuN^+^ cells in the lesion area and distal region. n.d., not detected. (N) Experiment timeline of TMEM119-ND1 mice, TBI was performed after tamoxifen induction. (O) Representative confocal images of GFP, IBA1 and NeuN in TMEM119-ND1 mice, showing no co-localization of GFP with NeuN in either the lesion core or distal regions. Scale bar, 50 μm. (P) Quantification of the percentage of GFP^+^NeuN^+^ cells among total NeuN^+^ cells in the lesion area and distal region. n.d., not detected.

Given tamoxifen was administrated after TBI induction in CX3CR1-ND1 mice, when circulating monocytes and macrophages were recruited in to injury region, the CX3CR1-CreER system can also label these peripheral myeloid cells during this period (29, 36) To exclude the recruited myeloid cells mis-led the lineage tracing result, we repeated the same experiments in TMEM119-ND1 mice (Figure 4K). The TMEM119-CreER system allows us to restrict NeuroD1 and GFP expression specifically to resident microglia in the brain (28). Consistent with the observations in CX3CR1-ND1 mice, we did not detect any GFP and NeuN co-localized cells in either the lesion core or regions distal to the injury site (Figure 4L). In addition, we reserved the sequency by conducting TBI after tamoxifen treatment, in turn, labeling resident microglia before the recruiting of peripheral myeloid cells (Figure 4N). Similarly, we observed no GFP-positive cells co-expressing pan neuron marker NeuN in this condition as well (Figure 4O). These findings further proved that NeuroD1 does not reprogram resident microglia into neurons even under injury environment.

## Discussion

In current study, we employed virus-free genetic lineage tracing system based on CX3CR1-creER and TMEM119-creER to trace the cell fate of NeuronD1 expressing cells, instead of viral based approaches. Through immunostaining assay at multiple timepoints, two-photon live cell imaging and scRNA sequencing, we systematically assessed the cell fate of NeuroD1-expressing microglia under both physiological and injury conditions. Collectively, all concordance observations provide strong evidence that NeuroD1-expressing microglia consistently retained their microglial identity and failed to acquire neuronal cell fate. These findings in current study contrast with previous reports claiming NeuroD1-mediated microglia-to-neuron conversion(24, 26), and instead support the notion that NeuroD1 is insufficient to induce microglia-to-neuron conversion. We speculated that the observed microglia to neuron conversion reported in previous study may have resulted from off-target viral expression, which leading to the mis interpretation of the cell source of the observed neurons (13, 21, 22, 25).

Previous studies have highlighted the controversial observations in validating direct glia-to-neuron reprogramming(13, 25). While several studies demonstrated that the single transcription factors, such as SOX2, ASCL1, or NeuroD1, could convert glial cells into neurons, subsequent investigations using rigorous lineage tracing or live imaging have failed to reproduce some of these inter-lineage conversions(20, 35). In fact, our longitudinal live cell imaging revealed a gradual loss of NeuroD1-expressing microglia over time. These findings were consitent with our previous results that ectopic NeuroD1 facilitated microglia cell apoptosis (13). The gradual disappearance of NeuroD1-expressing microglia overtime, LC3/P62 accumulation and increased TUNEL positivity, collectively indicate that NeuroD1 compromises microglial homeostasis and facilitate cell death rather than induces microglia-to-neuron conversion. Although the present study suggests that *in situ* microglia-to-neuron conversion remains technically challenging and require novel approaches, our team has developed an alternative therapeutic strategy, that is microglia replacement by bone marrow transplantation (Mr BMT) (37, 38). Mr BMT allows healthy microglia carrying normal or functional genes to replace diseased microglia carrying mutated genes, therefore providing an avenue to cure disease associated with primary microgliopathies (37). Indeed, Mr BMT has been successfully translated into clinical practice and was shown to halt disease progression in patients with adult-onset leukoencephalopathy with axonal spheroids and pigmented glia (ALSP), supporting Mr BMT as a promising cell-based therapies in neurological diseases(39).

Together, these findings underscore the importance of stringent lineage-tracing approaches and multimodal validation strategies when evaluating cell fate conversion. Although glia-to-neuron reprogramming remains a promising avenue for regenerative medicine, accumulating studies, including the present study raise the caution in interpreting in vivo glia-to-neuron conversion that without rigors validation of cellular origin and cell fate transition, particularly when considering the clinical translation of such approaches.

## Materials and Methods

### Animals

CX3CR1-CreER[B6.129P2(C)-Cx3cr1^tm2.1(cre/ERT2)Jung^/J, Stock No: 020940], Ai14 [B6.Cg-Gt(ROSA)26Sor^tm14(CAG-tdTomato)Hze^/J, Stock No: 007914], TMEM119-CreER (C57BL/6-Tmem119^em1(cre/ERT2)Gfng/J^, Stock No: 031820) and LSL-Neurod1-GFP [Gt(ROSA)26Sor^tm1(Neurod1-EGFP)Able^/BfriJ, Stock No: 034697] mice were purchased from The Jackson Laboratory. CX3CR1-CreER mice were crossed with LSL-Neurod1-EGFP mice to obtain CX3CR1-CreER^+/CreER^::LSL-Neurod1-EGFP^+/+^ (CX3CR1-ND1) mice. Similarly, CX3CR1-CreER mice were crossed with Ai14 mice to obtain CX3CR1-CreER::Ai14 (CX3CR1-Ai14) mice. All animals were housed in the Animal Facility of the Department of Laboratory Animal Science at Fudan University on a 12-h light‒dark cycle with food and water provided ad libitum. No significant sex differences were observed in this study.

### Drug administration

For mice aged 8 weeks or oder, CX3CR1-ND1, CX3CR1-Ai14, TMEM119-ND1 and TMEM119-Ai14 mice, tamoxifen at 150 mg/kg body weight was administered via intragastric gavage (i.g.) or intraperitoneal injection (i.p.) for four consecutive days to induce loxP-dependent recombination. For mice aged 4 to 6 weeks, the dose was reduced to 75 mg/kg body weight with the same adminstration sechdule.

### Tissue preparation for immunohistochemistry

The animals were deeply anesthetized with 1% pentobarbital sodium (60 mg/kg) via intraperitoneal injection and transcardially perfused with cold 0.01 M PBS and cold 4% paraformaldehyde (PFA). The brains were immediately separated from the skulls and postfixed in 4% PFA for 24h at 4°C. For vibratome sectioning, brains were directly cut into coronal sections at the desired thickness using a Leica vibratome after post-fixation. For cryo-sectioning, brains were dehydrated in 30% sucrose in 0.01 M PBS at 4 °C for 2–3 days, followed by immersion in fresh 30% sucrose for an additional day. Tissue sections containing the regions of interest were then cut at 35 or 40 μm thickness using a Leica CM1950 cryostat.

### Immunohistochemistry

The brain sections were rinsed 3 times in 0.01 M PBS for 10 min. The samples were blocked with 0.01 M PBS containing 0.3% Triton X-100 and 4% normal donkey serum (NDS, Jackson ImmunoResearch, Cat#: 017-000-121) at room temperature (RT) for 2 h. The samples were subsequently incubated with primary antibodies in 0.3% PBST containing 1% NDS at RT overnight. The samples were rinsed 3 times with 0.03% PBST for 10 min and then incubated with fluorescent dye-conjugated secondary antibodies in 0.03% PBST containing 1% NDS at RT for 2 h, together with 1:500 4’,6-diamidino-2-phenylindole (DAPI, Sigma‒Aldrich, Cat#: D9542-10MG). Finally, the samples were rinsed with 0.03% PBST for 10 min 3 times, and the slides were mounted with antifade mounting medium. The edges of the cover glass were then sealed with nail enamel.

In the present study, the following primary antibodies were used, including rabbit anti-Iba1 (Wako, Cat: 019-19741, 1:500), goat anti-IBA1 (Abcam, Cat: ab5076, 1:500), goat anti-GFP (Abcam, Cat: AB6673, 1:500), goat anti-mCherry (Biorbyt, Cat: orb11618, 1:500), rabbit anti-GFP (Invitrogen, Cat: A-11122, 1:1000), mouse anti-NEUN (Abcam, Cat: ab104224, 1:500), rabbit anti-NeuroD1 (Abcam, Cat: ab109224, 1:500), rabbit anti-RFP (Abcam, Cat: ab62341, 1:1000), rabbit anti-DCX (Abcam, Cat: ab18723, 1:200), mouse anti-MAP2 (Sigma, Cat: M9942, 1:2000), and rabbit anti-TUJ1 (Sigma, Cat: T3952, 1:2000), mouse anti-LC3B (Cell Signaling Technology, Cat: 83506T, 1:200), and rabbit anti-p62 (Cell Signaling Technology, Cat: 23214T, 1:500). The secondary antibodies included AF488 donkey anti-goat IgG (Jackson ImmunoResearch, Cat: 705-545-003, 1:2000), AF488 donkey anti-rabbit IgG (Jackson ImmunoResearch, Cat: 711-545-152, 1:2000), AF568 donkey anti-rabbit IgG (Invitrogen by Thermo Fisher Scientific, Cat: A10042, 1:2000), AF568 donkey anti-goat IgG (Invitrogen by Thermo Fisher Scientific, Cat: A11057, 1:2000), AF647 donkey anti-rabbit IgG (Jackson ImmunoResearch, Cat: 711-605-152, 1:2000), AF647 donkey anti-mouse IgG (Jackson ImmunoResearch, Cat: 715-605-151, 1:2000), and AF647 donkey anti-goat IgG (Jackson ImmunoResearch, Cat: 705-605-003, 1:2000).

### In situ apoptosis assay

A commercial Tunnel Cell Apoptosis Detection Kit (Service bio, Cat#: G1502-50T; Elab science, Cat#: E-CK-A321) was used to detect in situ cell apoptosis. Briefly, brain sections were treated with Proteinase K (20 ug/mL) for 10 mins at RT. Then, brain sections were incubated with equilibration buffer for 10 minutes at RT. Next, apoptotic cells were labeled by TMR-5-dUTP or FITC-dUTP labeling mix at RT and counterstained with DAPI. Positive controls were treated with DNase I and negative controls were labeled by buffer without recombinant TdT enzyme.

### Confocal microscopy

Confocal images of the fluorescent samples were acquired via 20x (NA 0.75), 40x (NA 1.30) or 60x (NA 1.40) objectives of a Nikon A2 confocal microscope. The Z-stacked focal planes were taken and maximally projected by ImageJ, whereas brightness and contrast were adjusted in ImageJ if necessary.

Tile scans of whole coronal brain sections were captured with an Olympus VS120 or VS200 system and exported from OlyVIA Software in TIFF format. Further data analysis was carried out in ImageJ.

### Two-photon surgery and imaging

Two-photon cranial window surgery was performed on mice anesthetized with isoflurane (3% induction, 1.2%-1.5% maintenance). A 3.5 mm-diameter circular cranial window was created at the skull above the sensory cortex, and a same-diameter circular D263T glass slice was implanted. The cranial window and a head bar were bonded using glue and dental cement to maintain stability. In addition to performing routine postoperative care, the mice were injected intraperitoneally with antibiotics and anti-inflammatory drugs (dexamethasone sodium phosphate injection and ceftiofur sodium for injection) for five consecutive days.

Two-photon imaging was performed on days 18, 22, 26, and 30 post-tamoxifen induction (as shown in the timeline in Fig. 2). During imaging, the mice were anesthetized with isoflurane (3% induction, 1.2%-1.5% maintenance), and body temperature was maintained by a heating pad. Imaging was performed via a self-developed two-photon microscopy system (developed by the Institute for Translational Brain Research, Fudan University) with one resonant and one galvo scan mirror. The excitation was delivered with a Coherent Chameleon Ti-Sapphire pulsed laser. The fluorescence emission was filtered with a Semrork FF01-520/35-25, FF01-593/46 filter and detected with a Hamamatsu photomultiplier tube (H7422-40P). The EGFP signal in Cx3cr1-CreER::LSL-Neurod1-EGFP mice was excited by a 920 nm laser and imaged via a 16x, 0.8 NA water-dipping objective (Nikon). Images were acquired at a depth of approximately 100 μm below the dura using ScanImage(40) at step intervals of 2 μm. Each Z-axis plane image (512x512 pixels) was imaged at a consistent zoom factor (∼300 × 300 μm area) and acquired at a 1.06 Hz frame rate.

Image processing was performed via ImageJ, and the Z-stack image containing the target cells was selected. Brightness and contrast were adjusted if needed, and the maximal projection was performed. Video processing was performed via Adobe Premiere software.

### AAV vectors, lentivirus preparation, and stereotaxic injection

The AAV vectors were purchased from Obio Technology, Ltd., and PackGene Biotech, Inc., and were as follows: *AAV1/2-CMV-EGFP, AAV5/2-CMV-EGFP*, *AAV6/2-CMV-EGFP*, *AAV8/2-CMV-EGFP*, *AAV9/2-CMV-EGFP*, *AAV2-DJ-CMV-EGFP*, *AAV2-retro-CMV-EGFP*, *AAV-PHP.eB-CMV-EGFP*, *AAV6-CX3CR1-GFP*, *AAV8-CX3CR1-GFP*, *AAV9-CX3CR1-GFP*, *AAV6-CD68-GFP*, *AAV8-CD68-GFP,* and *AAV9-CD68-GFP*.

All lentiviruses were packaged by Obio Technology, Ltd., according to the author’s design and requirements. All lentiviruses were packaged in HEK293 cells, and the packaged plasmids consisted of pMDL, VSV-G and pREV. The following lentiviral vectors were used: *pLenti-CAG-flag-NeuorD*1, *pLenti-CAG-flag-MCS*, *pLenti-CMV-GFP, pLenti-CX3CR1-GFP* and *pLenti-CD68-GFP*.

For intracranial injection, the mouse was anesthetized with isoflurane. The stereotaxic injection coordinates were +1.0 mm anterior/posterior, ± 1.7 mm medial/lateral, and -0.75 mm dorsal/ventral.

### Traumatic brain injury (TBI)

TBI was induced using the controlled cortical impact (CCI) model as previously described(36). Briefly, mice were anesthetized with ketamine (100 mg/kg) and xylazine (10 mg/kg) and placed in a stereotaxic frame (Stoelting, Wood Dale, IL, USA). Body temperature was maintained at 37.0 ± 0.5 °C using a heating pad throughout the procedure. After scalp disinfection, a midline incision (1.5–2 cm) was made to expose the skull. A 4 mm craniotomy was performed over the right parietal cortex (between bregma and lambda, 1 mm lateral to the midline) using a portable drill, taking care to preserve the dura mater. Mice with damaged dura were excluded from further analysis. Cortical impact was delivered using a 3 mm rounded steel impactor (PinPoint PCI3000, Hatteras Instruments Inc., USA) at a velocity of 1.5 m/s, impact depth of 1.5 mm, and dwell time of 100 ms. After impact, the bone flap was sealed with sterile bone wax and the scalp was sutured. Sham-operated mice underwent the same surgical procedures without cortical impact. Following surgery, mice were placed in a 37 °C recovery chamber until fully ambulatory.

### Isolation of brain CD11b^+^ cells by magnetic-activated cell sorting (MACS)

Briefly, mice were deeply anesthetized with isoflurane and transcardially perfused with cold normal saline. Brains were harvested, rinsed briefly in cold normal saline, and mechanically minced using a blade in a brain mold. The minced tissue was transferred into a C-tube containing 3 ml of digestion buffer per brain. The digestion buffer consisted of DMEM (Gibco, REF: C11995500BT) supplemented with papain (8 U/ml, Worthington, cat: LS00312), DNase I (125 U/ml, Sangon, cat: B100649-0040), and trypsin inhibitor (1.5 mg/ml, SIGMA, cat: T9253 or Aladdin, cat: A274384). Tissue dissociation was performed using a gentleMACS Octo Dissociator with Heaters (Miltenyi, 37°C_ABCK1 program). The resulting cell suspension was gently triturated, and the C-tube was rinsed with 0.5% BSA in DMEM. The suspension was passed through a 70-μm cell strainer (Falcon, cat: CLS431751) and collected in a 50-ml tube. Cells were centrifuged at 300 × g for 5 min at 4°C, and the pellet was resuspended in 4 ml of 30% Percoll (Cytiva, cat:17544502) prepared in 0.5% BSA in DMEM. Following centrifugation at 300 × g for 5 min at 4°C, the supernatant was carefully removed, and the cell pellet was washed once with 500 μl of 0.5% BSA in DPBS by centrifugation at 300 × g for 5 min at 4°C. The cell pellet was then resuspended in 90 μl of 0.5% BSA in DPBS and incubated with 10 μl of anti-CD11b magnetic beads (Miltenyi, cat: 130-097-142) for 30 min at 4°C. The labeled cells were loaded onto an LS Column (Miltenyi, cat: 130-042-401) pre-equilibrated with 10 ml of 0.5% BSA in DPBS and washed with an additional 10 ml of the same buffer. The column was subsequently removed from the magnetic separator, and the magnetically labeled CD11b^+^ cells were eluted with 5 ml of 0.5% BSA in DPBS by firmly applying the plunger twice. The eluted cells were collected and centrifuged at 300 × g for 5 min at 4°C, followed by resuspension in the appropriate buffer for cell counting and downstream experiments.

### Rotarod test

All mice were acclimated to the behavioral room for half an hour before test. Briefly, in the training phase, the mice were placed on rotating rod at 5 revolution per minute (RPM) to learn how to walk forward and maintain balance. Each mouse was placed on the rod for 60 seconds and then returned back to its home cage. This procedure was repeated three times with 10 minuties intervals. In the testing phase, mice were placed in separate lanes on the rod, which was set to accelerate from 4 to 40 RPM over a 300-second. This test was repeated three times, with approximately 30 minutes intervals between trails. The latency to fall was recorded and used for subsequent statistical analysis.

### Grip strength test

All mice were acclimate to the behavioral room for half an hour before test. In brief, we gently held mouse by the tail and placed it with all four paws on the grip strength grid. The mouse was then gently pulled back until it relased the grasp, and the holding force was recorded. This mearsurement was repeated six times per mouse, and the maximum force value was used for subsequent statistical analysis.

### Single-cell preparation and fluorescence-activated cell sorting (FACS)

For scRNA-seq, FACS was used to harvest GFP^+^ or tdTomato^+^ cells from tamoxifen-treated *Cx3cr1*-*creER*::*LSL-Neurod1-GFP* and *Cx3cr1-CreER*::*Ai14* mice. Briefly, brains were dissected and cut into 1-mm³ pieces using a mouse stainless steel brain matrix (RWD). The tissue pieces were digested in 3 ml of DMEM containing 8 U/ml papain (8 U/ml, Worthington, cat: LS00312), 125 U/ml DNase I (Sangon Biotech, cat: B100649-0040), and 1.5 mg/ml trypsin inhibitor (SIGMA, cat: T9253 or Aladdin, cat: A274384) via a gentleMACS Octo Dissociator (Miltenyi) with the 37C_ABDK program.

After 30 min of digestion, the mixture was gently pipetted up and down and then filtered through a 70 µm cell strainer. The dissociated cells were centrifuged at 370 × g at RT for 10 min, and the supernatant was discarded. The cell pellets were resuspended in 4 ml of 30% Percoll (Cytiva, cat: 17089101) and centrifuged again at 370 × g and 17°C for 10 min to remove debris. After centrifugation, the supernatant was discarded, and the cells were washed once with DPBS containing 0.5% BSA. Before the cells were loaded onto the SONY MA900, they were stained with 7-AAD (BD Pharmingen, cat: 559925, 1:700) to label dead cells. Finally, the tdTomato^+^ 7-AAD^-^ or GFP^+^ 7-AAD^-^ cells were collected for scRNA-seq immediately.

### scRNA-seq data analysis

scRNA-seq was conducted via the MobiCube High-throughput Single Cell 3 ′ Transcriptome Set V2.1 (PN-S050200301) and the MobiNova-100 microfluidic platform. The single-cell suspension was adjusted to a concentration of 700–1200 cells/μl and immediately loaded onto a chip for microdroplet formation via the MobiNova-100 system. Reverse transcription, cDNA amplification, and DNA library construction were carried out according to the manufacturer’s protocol. High-throughput sequencing was performed in PE-150 mode.

FASTQ files were processed and aligned to the mouse reference genome (mm10) via MobiVision version 3.2, with unique molecular identifier (UMI) counts aggregated for each barcode. The resulting UMI count matrix was analyzed in R version 4.2.2 via Seurat version 4.3.0. To ensure data quality, we excluded low-quality cells and potential multiplets on the basis of the following criteria: cells with fewer than 200 genes or more than 5000 genes and cells with more than 10% UMIs mapped to mitochondrial genes. scRNA-seq data from the four groups were integrated via *IntegratedData* functions according to the manufacturer’s instructions. The remaining data were normalized via the *NormalizeData* function and scaled with *ScaleData*. For scRNA-seq analysis, principal component analysis (PCA) was conducted on variable features, with the top 30 principal components (PCs) selected for clustering and visualization via tSNE and uniform manifold approximation and projection (UMAP). Cell clustering was performed with a resolution of 1 to annotate the cell types. Differentially expressed genes (DEGs) among cell groups were identified via the *FindMarkers* function with parameters set to min.pct = 0.1 and a log fold change (FC) threshold of 0.25 (Wilcoxon rank-sum test). Genes with |log2FC| > 0.25 and P < 0.05 were considered significant DEGs and subsequently used for Gene Ontology (GO) analysis. Visualization of the results was performed via the *VlnPlot* and *FeaturePlot* functions. We gratefully thank OE Biotech Co., Ltd. (Shanghai, China) for providing sequencing service.

### Statistical analysis

Statistical analyses were performed with Prism 10.2.0 (GraphPad). Each data point represents the average statistical result of more than three brain sections in different bregma regions of one mouse. The results were evaluated independently in a double-blind manner. All the results are presented as the means ± standard deviations (SDs). One-way analysis of variance (ANOVA) with Holm‒Sidak’s multiple comparisons test (post hoc) was performed for multiple comparisons, whereas a two-tailed independent t test was used to compare the differences between two groups. No outliers were excluded.

## Data availability

The sequencing data generated in this study have been deposited in the Gene Expression Omnibus under accession number GSE277222. All other data supporting the findings of this study are available from the corresponding author upon reasonable request.

## Author Contributions

Y.R. and B.P. conceived and designed this study. Y.R. and B.P. supervised and conceptualized this study. Y.R., and X.L. wrote the manuscript. X.L., Y.L., Y.C., N.H., P.O., Y.J., and S.G. performed most experiments. X.L. performed data analysis. L. C. provided necessary study support. All authors discussed the results and commented on this manuscript.

## Acknowledgments

This work was supported by Brain Science and Brain-like Intelligence Technology–National Science and Technology Major Project (2022ZD0207200, R.Y., 2022ZD0204700, P.B.); National Natural Science Foundation of China (82621102, P.B., 32571128, R.Y., 323B2030, L.X., 325B2039, O.P.); the Fellowship of China National Postdoctoral Program for Innovative Talents (BX20250132, L.X.); the Fellowship of China Postdoctoral Science Foundation (2025M782569, L.X.); Shanghai Pilot Program for Basic Research (21TQ014, P.B.), Changping Laboratory (2025B-07-18, P.B.) and Lin Gang Laboratory (LGL-8998-02, P.B.). In addition, authors also express their gratitude and respect to all animals sacrificed in this study. During the preparation of this manuscript, the authors, as the non-native English speakers, used ChatGPT 4o and 5 to improve the language and enhance its readability.

## Declaration of generative AI and AI-assisted technologies in the manuscript preparation process

As the authors are non-native English speakers, the authors used ChatGPT 5.5 during the preparation of this manuscript in order to improve the language and enhance its readability. After using this tool, the authors reviewed and edited the content as needed and take full responsibility for the content of the published article.

## Supporting Information

**Movie 1.** This video shows that GFP^+^ cells did not convert to neurons in vivo via 2-photon imaging at 18, 22, 26 and 30 days post-tamoxifen administration. Refers to Figure 3.

**Fig. S1.**
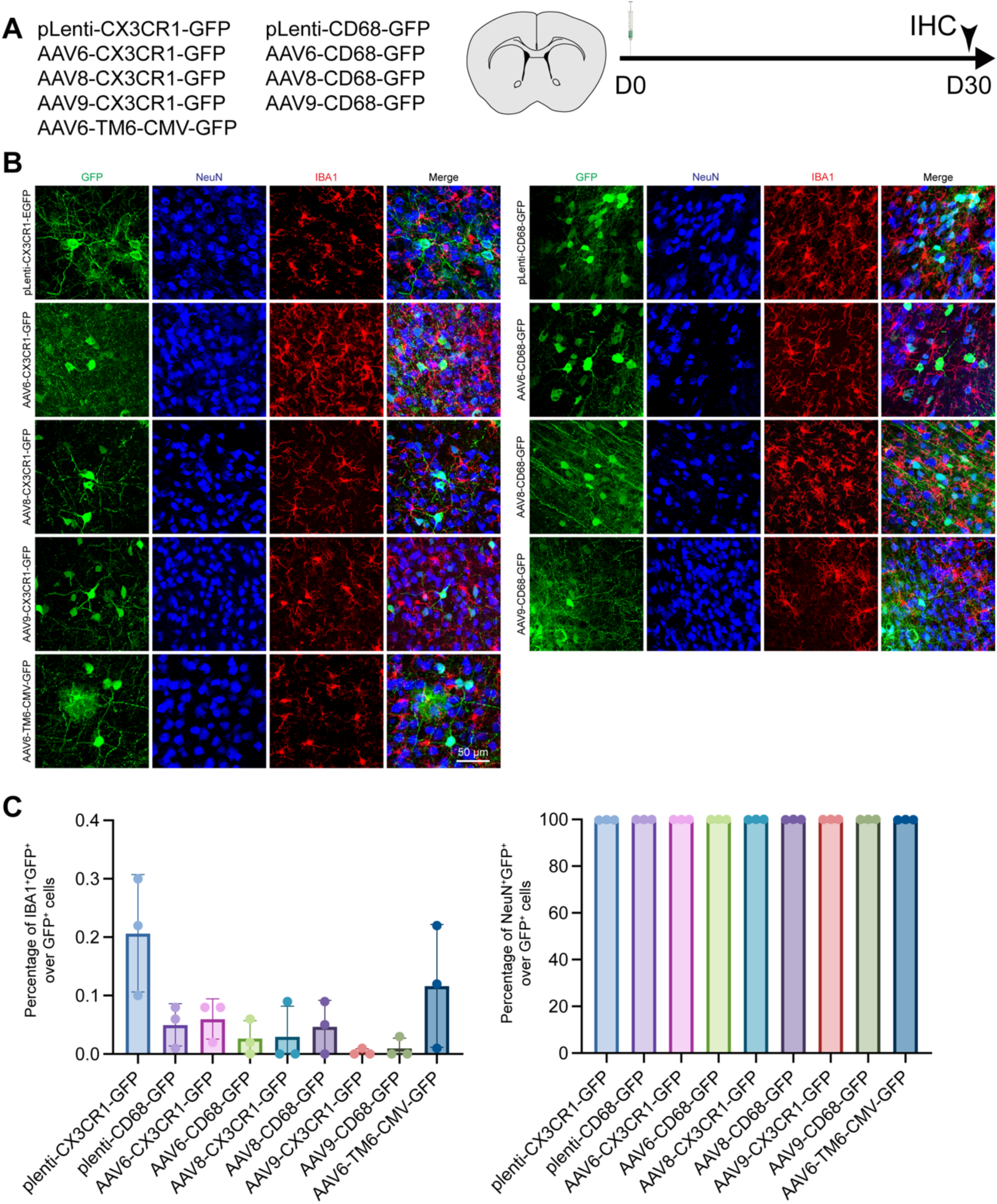
Lentivirus- or AAV-based infection caused leaky neuronal expression *in vivo*. (A) The study design shows the different viruses used here. (B) Representative confocal images showing the expression of GFP, IBA1 and NEUN in the brains of the lentivirus- or AAV-infected mice. (C) The efficacy of infection was quantified, and the data are presented as the ratio of IBA1^+^GFP^+^/GFP^+^ cells. N = 3 mice for each group.

**Fig. S2.**
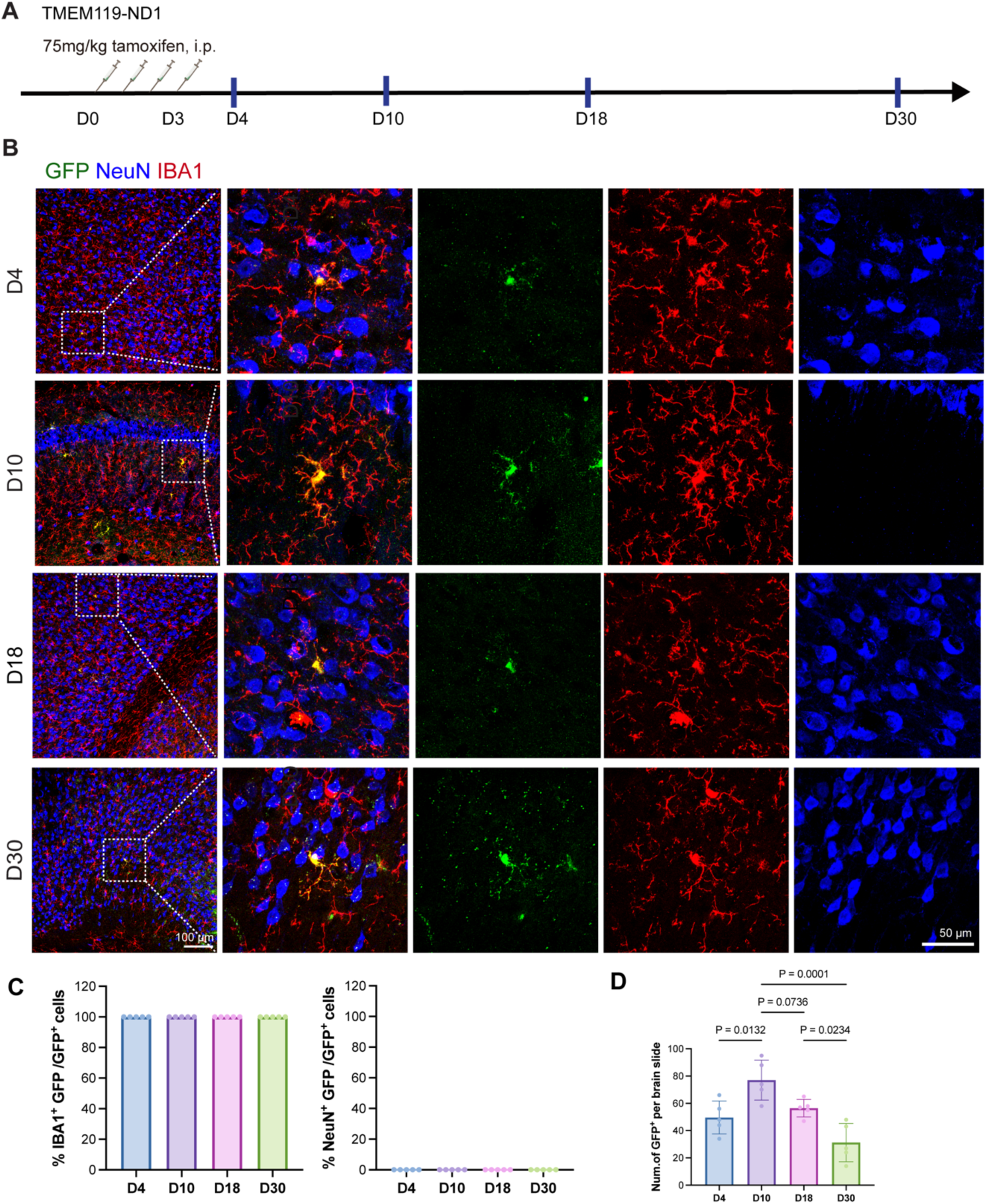
NeuroD1-expressing microglia is co-labeled with microglial marker IBA1 but not neuronal marker NeuN. (A) Experimental timeline. (B) Representative confocal images at D4, D10, D18 and D30 showing that GFP is robustly co-expressed in IBA1^+^ cells but not NeuN^+^ cells after Cre recombination. (C) Quantification of IBA1^+^GFP^+^ double-positive cells in GFP^+^ cells and NeuN^+^GFP^+^ in GFP^+^ cells at D4, D10, D18 and D30. N = 5 mice for each group. (D) Quantification of GFP^+^ cells per brain slide in TMEM119-ND1 mice. N = 5 mice for each group.

**Fig. S3.**
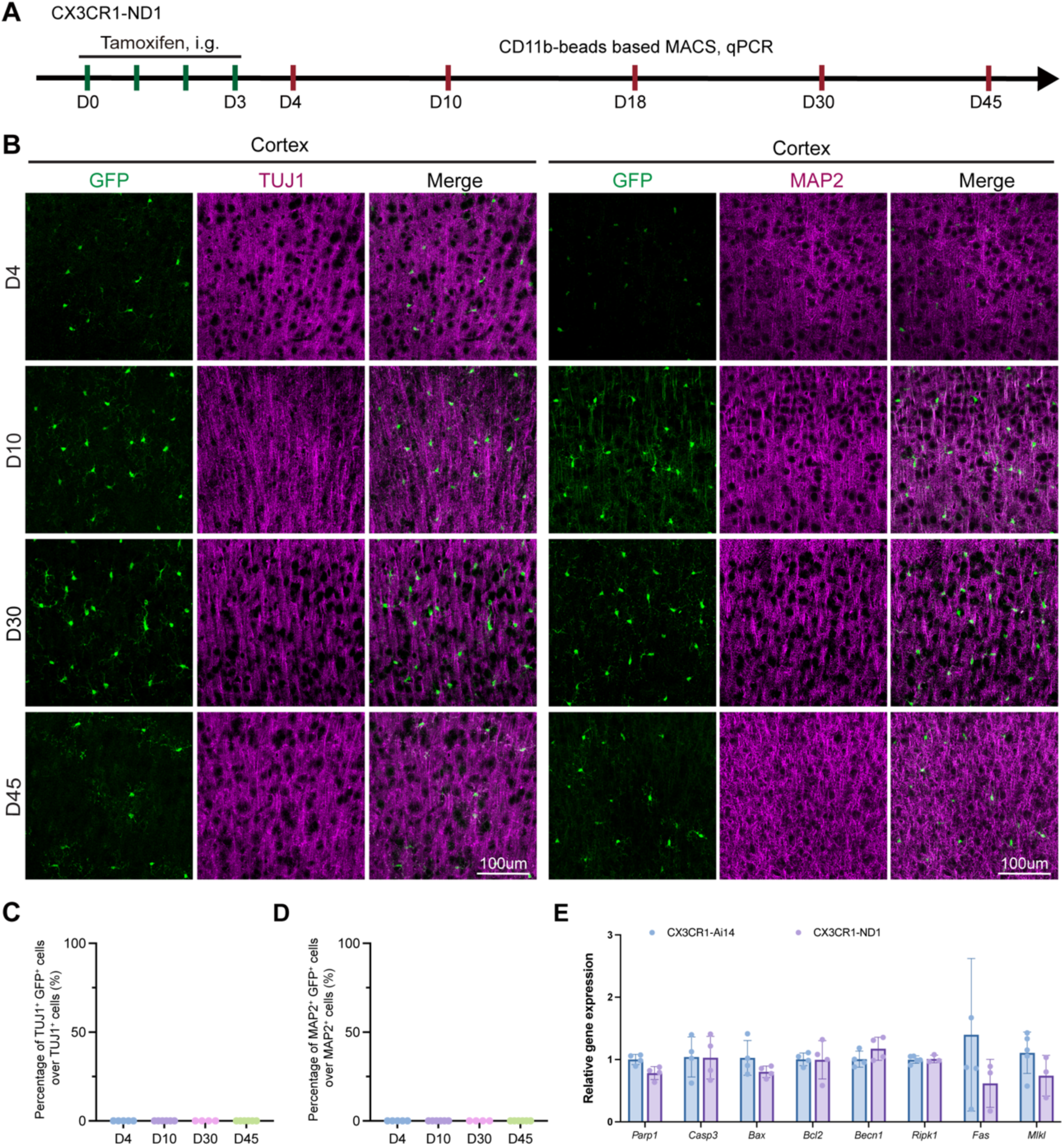
Ectopic NeuroD1 expression in microglia does not induce microglia-to-neuron conversion, as GFP does not co-localize with TUJ1 and MAP2. (A) Experimental timeline. (B) Representative confocal images at D4, D10, D30 and D45 showing that GFP is not co-labeled with TUJ1 or MAP2 in the brain. (C) Quantification of TUJ1^+^GFP^+^ double-positive cells among TUJ1^+^ cells at D4, D10, D30 and D45. N = 4 or 5 mice for each group. (D) Quantification of MAP2^+^GFP^+^ double-positive cells among MAP2^+^ cells at D4, D10, D30 and D45. N = 4 or 5 mice for each group. (E) Relative gene expression level at D18. N = 4 mice for each group.

**Fig. S4.**
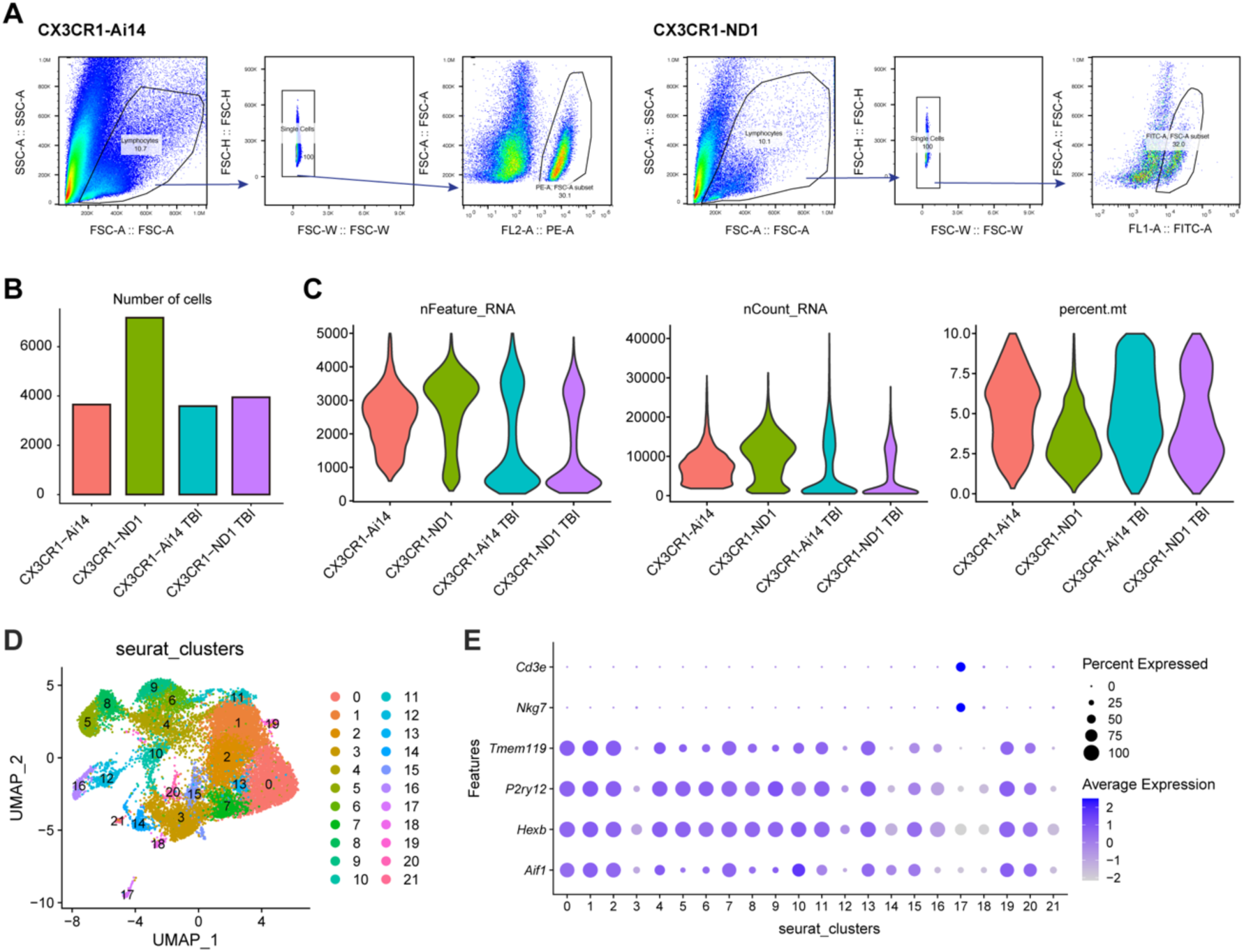
FACS gating strategy and supplemental quality control information of scRNA-seq data. (A) Gating strategy of CX3CR1-Ai14 and CX3CR1-ND1 mice. (B) Number of cells (after quality control) per group. (C) Violin plot shows the number of genes detected in each cell (nFeature_RNA), total number of molecules detected within a cell (nCount_RNA) and the distribution of mitochondrial genes in each cell (percent.mt) among the 4 groups. (D) UMAP plot show all cells were divided into 22 unsupervised cluster with resolution = 1. (E) Dotplot show microglia, T cell and NK cell specific markers expression over 22 clusters.

## References

1. N. H. Varvel et al., Microglial repopulation model reveals a robust homeostatic process for replacing CNS myeloid cells. Proc Natl Acad Sci U S A 109, 18150–18155 (2012).

2. Y. Huang et al., Repopulated microglia are solely derived from the proliferation of residual microglia after acute depletion. Nature neuroscience 21, 530–540 (2018).

3. X. Li et al., Transcriptional and epigenetic decoding of the microglial aging process. Nat Aging 10.1038/s43587-023-00479-x, 1288–1311 (2023).

4. Z. Guo et al., In vivo direct reprogramming of reactive glial cells into functional neurons after brain injury and in an Alzheimer’s disease model. Cell stem cell 14, 188–202 (2014).

5. N. Heins et al., Glial cells generate neurons: the role of the transcription factor Pax6. Nature neuroscience 5, 308–315 (2002).

6. W. Tai et al., In vivo reprogramming of NG2 glia enables adult neurogenesis and functional recovery following spinal cord injury. Cell Stem Cell 28, 923–937.e924 (2021).

7. W. Niu et al., In vivo reprogramming of astrocytes to neuroblasts in the adult brain. Nature Cell Biology 15, 1164–1175 (2013).

8. W. Niu et al., SOX2 reprograms resident astrocytes into neural progenitors in the adult brain. Stem Cell Reports 4, 780–794 (2015).

9. S. Chanda et al., Generation of induced neuronal cells by the single reprogramming factor ASCL1. Stem Cell Reports 3, 282–296 (2014).

10. H. Qian et al., Reversing a model of Parkinson’s disease with in situ converted nigral neurons. Nature 582, 550–556 (2020).

11. H. Zhou et al., Glia-to-Neuron Conversion by CRISPR-CasRx Alleviates Symptoms of Neurological Disease in Mice. Cell 10.1016/j.cell.2020.03.024 (2020).

12. C. N. Svendsen, M. V. Sofroniew, Lineage tracing: The gold standard to claim direct reprogramming in vivo. Mol Ther 30, 988–989 (2022).

13. Y. Rao et al., NeuroD1 induces microglial apoptosis and cannot induce microglia-to-neuron cross-lineage reprogramming. Neuron 109, 4094–4108 e4095 (2021).

14. T. Hoang et al., Ptbp1 deletion does not induce astrocyte-to-neuron conversion. Nature 618, E1–E7 (2023).

15. G. Yang et al., Ptbp1 knockdown failed to induce astrocytes to neurons in vivo. Gene Ther 30, 801–806 (2023).

16. Y. Xie, J. Zhou, B. Chen, Critical examination of Ptbp1-mediated glia-to-neuron conversion in the mouse retina. Cell reports 39, 110960 (2022).

17. T. Guo et al., Downregulating PTBP1 fails to convert astrocytes into hippocampal neurons and to alleviate symptoms in Alzheimer’s mouse models. J Neurosci 10.1523/jneurosci.1060-22.2022 (2022).

18. T. Hoang et al., Genetic loss of function of Ptbp1 does not induce glia-to-neuron conversion in retina. Cell reports 39 (2022).

19. W. Chen, Q. Zheng, Q. Huang, S. Ma, M. Li, Repressing PTBP1 fails to convert reactive astrocytes to dopaminergic neurons in a 6-hydroxydopamine mouse model of Parkinson’s disease. eLife 11, e75636 (2022).

20. Y. Xie, J. Zhou, L. L. Wang, C. L. Zhang, B. Chen, New AAV tools fail to detect Neurod1-mediated neuronal conversion of Muller glia and astrocytes in vivo. EBioMedicine 90, 104531 (2023).

21. D. Leib, Y. H. Chen, A. M. Monteys, B. L. Davidson, Limited astrocyte-to-neuron conversion in the mouse brain using NeuroD1 overexpression. Mol Ther 30, 982–986 (2022).

22. L. L. Wang et al., Revisiting astrocyte to neuron conversion with lineage tracing in vivo. Cell 184, 5465–5481.e5416 (2021).

23. T. Matsuda et al., Pioneer Factor NeuroD1 Rearranges Transcriptional and Epigenetic Profiles to Execute Microglia-Neuron Conversion. Neuron 101, 472–485 e477 (2019).

24. T. Matsuda, K. Nakashima, Clarifying the ability of NeuroD1 to convert mouse microglia into neurons. Neuron 10.1016/j.neuron.2021.11.012 (2021).

25. Y. Rao, B. Peng, Failure of observing NeuroD1-induced microglia-to-neuron conversion in vitro is not attributed to the low NeuroD1 expression level. Mol Brain 15, 31 (2022).

26. T. Irie et al., Direct neuronal conversion of microglia/macrophages reinstates neurological function after stroke. Proc Natl Acad Sci U S A 120, e2307972120 (2023).

27. A. M. Rosario et al., Microglia-specific targeting by novel capsid-modified AAV6 vectors. Mol Ther Methods Clin Dev 3, 16026 (2016).

28. A. M. Bedolla et al., A comparative evaluation of the strengths and potential caveats of the microglial inducible CreER mouse models. Cell Rep 43, 113660 (2024).

29. S. Yona et al., Fate Mapping Reveals Origins and Dynamics of Monocytes and Tissue Macrophages under Homeostasis. Immunity 38, 1073–1079 (2013).

30. H. J. Li et al., Intestinal Neurod1 expression impairs paneth cell differentiation and promotes enteroendocrine lineage specification. Sci Rep 9, 19489 (2019).

31. X. Guo, C. Wang, Y. Zhang, R. Wei, R. Xi, Cell-fate conversion of intestinal cells in adult Drosophila midgut by depleting a single transcription factor. Nat Commun 15, 2656 (2024).

32. M. Noguchi et al., Autophagy as a modulator of cell death machinery. Cell Death Dis 11, 517 (2020).

33. Q. Chen, J. Kang, C. Fu, The independence of and associations among apoptosis, autophagy, and necrosis. Signal Transduction and Targeted Therapy 3, 18 (2018).

34. H. Changotra et al., ATG5: A central autophagy regulator implicated in various human diseases. Cell Biochemistry and Function 40, 650–667 (2022).

35. C. Heinrich et al., Sox2-mediated conversion of NG2 glia into induced neurons in the injured adult cerebral cortex. Stem Cell Reports 3, 1000–1014 (2014).

36. L. Cai et al., ALKBH5 demethylates the m(6)A modification of SOCS3 in microglia/macrophages and alleviates neuroinflammation after brain injury. Proc Natl Acad Sci U S A 122, e2504697122 (2025).

37. Y. Rao, Y. Bai, X. Li, B. Du, B. Peng, The evolution of microglia replacement: A new paradigm for CNS disease therapy. Cell Stem Cell 32, 1807–1832 (2025).

38. Z. Xu et al., Efficient Strategies for Microglia Replacement in the Central Nervous System. Cell Rep 32, 108041 (2020).

39. J. Wu et al., Microglia replacement halts the progression of microgliopathy in mice and humans. Science 389, eadr1015 (2025).

40. T. A. Pologruto, B. L. Sabatini, K. Svoboda, ScanImage: Flexible software for operating laser scanning microscopes. BioMedical Engineering OnLine 2, 13 (2003).

